# Characterization of Identified Dopaminergic Neurons in the Mouse Forebrain and Midbrain

**DOI:** 10.1101/2023.08.29.554772

**Authors:** Maggy Yu Hei Lau, Sana Gadiwalla, Susan Jones, Elisa Galliano

## Abstract

Dopaminergic (DA) neurons play pivotal roles in diverse brain functions, spanning movement, reward processing, and sensory perception. DA neurons are most abundant in the midbrain (Substantia Nigra pars compacta, SNC, and Ventral Tegmental Area, VTA) and the olfactory bulb (OB) in the forebrain. Interestingly, a subtype of OB DA neurons is capable of regenerating throughout life, while a second class is exclusively born during embryonic development. Emerging evidence in SNC and VTA also indicates substantial heterogeneity in terms of morphology, connectivity, and function. To further investigate this heterogeneity and directly compare form and function of midbrain and forebrain DA neurons, we performed immunohistochemistry and whole-cell patch-clamp recordings in *ex vivo* brain slices from juvenile DAT-tdTomato mice. After confirming the penetrance and specificity of the dopamine transporter (DAT) Cre line, we compared soma shape, passive membrane properties, voltage sags and action potential firing across midbrain and forebrain DA subtypes. We found that each DA subgroup within midbrain and forebrain was highly heterogeneous, and that DA neurons across the two brain areas are also substantially different. These findings complement previous work in rats as well as gene expression and *in vivo* datasets, further questioning the existence of a single “dopaminergic” neuronal phenotype.

## Introduction

The neurotransmitter dopamine (DA) is instrumental in modulating wide-ranging functions, which include motor control, learning, and emotional regulation (Berke 2018). Given such broad behavioural reach, it is somewhat surprising that in the mammalian brain DA is only produced by a few clusters of cells which together account for less than one percent of the total neuron number(Chinta and Andersen 2005). Almost all dopaminergic (DA) neurons are localised to the midbrain cell groups A8 (retrorubal), A9 (substantia nigra pars compacta, SNC) and A10 (ventral tegmental area, VTA) (Dahlstroem and Fuxe 1964; Björklund and Dunnett 2007; Garritsen et al. 2023). These midbrain DA neurons have distinct inputs, axonal projections, functional roles, and clinical relevance (Ungless and Grace 2012). Malfunctioning, mis-development, or degeneration of these neurons results in pathologies such as Parkinson’s disease (PD) (Sillitoe and Vogel 2008; Poewe et al. 2017) (Poewe et al. 2017), while substance abuse disorders at least initially involve VTA projections to the ventral striatum (Lüscher, Robbins, and Everitt 2020). Thus unsurprisingly, the physiology and pathology of midbrain DA neurons have long been the focus of intense fundamental and translational research (Bissonette and Roesch 2016). Outside the midbrain resides the less investigated minority (10%) of DA groups. These are found in the retina, diencephalon (hypothalamic groups A11-15), and in the forebrain where group A16 in the olfactory bulb (OB) is the most abundant (Andén et al. 1965; Halász et al. 1981; Björklund and Dunnett 2007). OB DA neurons are local inhibitory interneurons which, by co-releasing DA and GABA in the bulbar glomerular layer, modulate the information transfer at the first synapse in the olfactory pathway and contribute to shape olfactory behaviour (Pignatelli et al. 2005; Hsia, Vincent, and Lledo 1999; Banerjee et al. 2015; Economo, Hansen, and Wachowiak 2016; McGann 2013). Interestingly, most OB DA neurons can regenerate throughout the animal’s lifetime (Bonzano et al. 2016). Indeed, an increase in their number, rather than the degeneration occurring in the midbrain, has been reported in PD patients, and given their inhibitory role it could account for the hyposmia/anosmia associated with the disease (Huisman, Uylings, and Hoogland 2004).

Within each anatomically-defined group, DA neurons have historically been considered homogeneous populations of cells which express tyrosine hydroxylase (TH, the rate limiting enzyme in the catecholamine synthesis (White and Thomas 2012). However, evidence for functional heterogeneity started to accumulate in the early 2000s, with the caveat that it largely relied on uncertain methods of DA identification, notably the action potential waveform in current clamp recordings, or the presence of a hyperpolarisation-activated cation current, Ih, in voltage clamp recordings (Margolis et al. 2006; Neuhoff et al. 2002; Pignatelli et al. 2005). The emergence of technologies allowing the generation of mouse lines which target specific cell types confirmed that DA neurons within the SNC, VTA and OB are highly heterogeneous in terms of their developmental profile, gene expression, morphology, electrophysiological properties, and/or connectivity (Roeper 2013; Morales and Margolis 2017; Kosaka, Pignatelli, and Kosaka 2020; Galliano et al. 2018; Greene 2006; Tiklová et al. 2019; Poulin et al. 2018; 2020; Azcorra et al. 2023; Brown et al. 2009; Lammel et al. 2015). Somewhat surprisingly, no study has yet compared the functional heterogeneity of genetically-identified DA neuron electrophysiological properties within VTA, and across the midbrain. To date, bulbar DA neurons show perhaps the highest level of morphological and functional heterogeneity and can be cleanly divided into two subgroups. While the smallest subgroup consists of neurons with far-branching dendrites and an axon, and which are born exclusively during early embryonic development (“standard” neurons, sometimes called short-axon cells), the majority of bulbar DA cells do not have an axon but release GABA and dopamine from narrowly-branched dendrites, and undergo life-long neurogenesis (Galliano et al. 2018; Kosaka, Pignatelli, and Kosaka 2020; Kiyokage et al. 2010; Bonzano et al. 2016; Chand et al. 2015; Tufo et al. 2022). The neurogenic properties of OB DA neurons and their ability to insert themselves into a pre-existing circuit have prompted their exploration as potential candidates for cell replacement therapies in pathologies marked by the degeneration of midbrain DA neurons (Deleidi et al. 2011; Cave, Wang, and Baker 2014). However, given the recent confirmation that only the anaxonic OB DA subtype can undergo adult neurogenesis, it is important to clarify whether they share any functional properties with the neurons that they could potentially replace.

In this study we took advantage of the DAT-Cre transgenic mouse line to insert a fluorescent tag in neurons expressing the dopamine transporter (DAT) promotor (Bäckman et al. 2006a). In thus identified DA neurons we investigated their morphological and functional heterogeneity within and between midbrain and forebrain. Overall, we aimed to answer this question: do all DA neurons share common neurophysiological features?

## Materials and Methods

### Animals

DAT-Cre (B6.SJL-Slc6a3 tm1.1(cre)Bkmn/J, Jax stock 006660) mice were crossed with flox-tdTomato (B6;129S6-Gt(ROSA)26Sortm9(CAG-tdTomato)Hze/J, Jax stock 007909). Transgenic DAT-Cre x flex-tdTomato (tdT) mice of either gender was used aged postnatal day 16-36 in all experiments in accordance with the Animals (Scientific procedures) Act 1986 and with AWERB (Animal Welfare and Ethical Review Board) approval. Mice were housed in 12-hour light-dark cycle in an environmentally controlled room with ad libitum access to water and food.

### Immunohistochemistry

Mice were anesthetized with an overdose of pentobarbital and then perfused with 20 mL PBS with heparin (20 units.mL^-1^), followed by 20mL of 1% paraformaldehyde (PFA; TAAB Laboratories; in 3% sucrose, 60 mM PIPES, 25 mM HEPES, 5 mM EGTA, and 1 mM MgCl_2_). The brains were dissected and post-fixed in 1% PFA for 2-7 days, then embedded in 5% agarose and sliced horizontally at 50μm using a vibratome (VT1000S, Leica). Free-floating slices containing SNC, VTA and OB were washed with PBS and incubated in 5% normal goat serum (NGS) in PBS/Triton/azide (0.25% Triton, 0.02% azide) for 2 h at room temperature. They were then incubated in primary antibody (tyrosine hydroxylase (TH), mouse, Millipore, 1:500; DsRed, rabbit, 1:500, Clontech) diluted in a PBS/Triton/azide for 2 days at 4°C. Slices were then washed three times for 5 min with PBS, before being incubated in secondary antibody solution (mouse 488 and rabbit 633 Alexa Fluor®; 1:1000 in PBS/Triton/azide) for 3h at room temperature. After washing in PBS, slices underwent an additional counterstaining step with 0.2% Sudan black in 70% ethanol at room temperature for 3 min to minimize autofluorescence, and then they were mounted on glass slides Menzel-Gläser) with FluorSave. Unless stated otherwise all reagents were purchased from Sigma.

### Fixed-tissue imaging and analysis

All images were acquired with a laser scanning confocal microscope (Zeiss LSM 710) using appropriate excitation and emission filters, a pinhole of 1 AU and a 40x oil immersion objective. Laser power and gain were set to prevent signal saturation. All quantitative analysis was performed with Fiji (Image J). Images were taken with a 0.7x zoom (0.593μm/pixel), 512x512 pixels, and in z-stacks with 1μm steps. In all animals, images were sampled from three slices taken from the upper, middle, and lower third of the portion of the brain in the dorsal-ventral axis containing the midbrain. SNC was defined as being adjacent (anteriorly and posteriorly) to the medial terminal nucleus of the accessory optic tract (MT); VTA was defined as being between MT and the midline. The OB was sampled both rostrally and caudally. To avoid selection biases, all cells present in the stack and positive for either identifying marker (TH or DsRed) were measured. DsRed immunofluorescence in the 633 channel was analysed instead of the genetically-encoded tdT fluorescence in the 580 channel to standardise comparison with the TH immunofluorescence and account for antibody penetration. TH and/or DsRed positive bulbar DA cells were included in the analysis only if their soma was in or bordering with the glomerular layer. For both the TH and DsRed channels the staining intensity of each ROI (expressed as mean grey value) was compared to the background staining in each image, defined by manually drawing a “background ROI” in an area clearly devoid of cells. A cell was considered TH- or DsRed-positive if its mean grey value in the respective channels was bigger than 1.2 times the background-ROI value. Soma area was measured at the single plane including the cell’s maximum diameter by drawing a region of interest (ROI) with the free-hand drawing tool. Fiji(ImageJ) shape descriptors analysis was used on these ROIs to extract maximum diameter (Feret’s diameter, the longest distance between any two points along the selection boundary), and roundness (4*area/(*majoraxis^2), the inverse of the aspect ratio). DA cell density was calculated for each image by dividing the number of analysed DsRed-positive cells by the volume of the SNC/VTA/glomerular layer (z depth x region of interest area, drawn and measured in a maximum intensity projection of the DsRed channel).

### Electrophysiology

Mice were anesthetized with isoflourane and then decapitated. The desired brain area (either midbrain or olfactory bulb) was then dissected and submerged in ice-cold sucrose solution containing (in mM): 240 sucrose, 5 KCl, 1.25 Na_2_HPO_4_, 2 MgSO_4_, 1 CaCl_2_, 26 NaHCO_3_ and 10 D-Glucose continuously bubbled with 95% O2 and 5% CO2. Horizontal slices of the midbrain (200-250μm) and olfactory bulb (300μm) were made using a Campden Instruments 7000 Vibroslicer. Slices were then transferred to a submersion incubation chamber with a glucose artificial cerebrospinal fluid (ACSF) solution containing (in mM): 124 NaCl, 5 KCl, 1.25 Na_2_HPO_4_, 5 MgSO_4_, 2 CaCl_2_, 26 NaHCO_3_ and 20 D-Glucose, maintained in 95% O2 and 5% CO2 for 30 minutes to 5 hours before patch clamp recordings.

Patch pipettes were made from borosilicate glass (PG150T-7.5, Harvard Apparatus Ltd., Kent, UK) to achieve a tip resistance of 1.5-5.0MOhm (larger tips for midbrain, smaller for OB) when filled with solution containing (in mM): 124 K-Gluconate, 9 KCl, 10 KOH, 4 NaCl, 10 HEPES, 28.5 Sucrose, 4 Na_2_ATP, 0.3 Na_3_GTP (pH 7.20 – 7.30; 270 – 300mOsm). DA cells were visualised using an upright microscope (Olympus BX51WI) equipped with a 4x air objective and a 40X water immersion objective, and SciCam camera (Scientifica). tdT fluorescence was revealed by LED (CoolLED pE-100) excitation. Under low power magnification, the glomerular layers in OB slices were visually located as circular structures proximal to the olfactory sensory nerves. The SNC and VTA in the midbrain slices were located by visually locating the medial terminal nucleus (MT) of the accessory optic tract. The SNC was identified as the region of DA neurons adjacent to MT and the VTA was identified as the region of DA neurons medial to the MT. During recordings slices were perfused with ACSF at 30±2°C with an in-line and chamber heater (Warner Instrument Corporation, MA, USA). Whole-cell patch clamp recordings were performed using an Axon Axopatch 200B patch clamp amplifier (Molecular Devices, CA, USA). Signals were filtered at 2kHz then acquired and digitized at 20 kHz using a Micro 1401 acquisition unit (Cambridge Electronic Design, Cambridge, UK). DA neurons were voltage clamped to -60mV and their passive properties were probed between each experimental protocol with test pulses of 5mV for 10ms. Recordings were excluded if the series resistance (Rs) exceeded 20MOhm (for midbrain neurons) or 30MOhm (for OB neurons); or input resistance (Ri) was lower than 100MOhm (both midbrain and OB neurons). Recordings were also excluded when two test pulses measured before and after the recording varied by over 20%. Experimental recordings were acquired and analysed using Spike 2 (Version 4; Cambridge Electronic Design, Cambridge, UK).

Single action potential (AP) experiments were performed in current clamp in continuous mode. DA neurons were clamped to -60 ± 3mV and 12 pulses of 10ms current of increasing amplitudes were injected at 3s intervals into midbrain DA neurons in 40pA increments (0 – 480pA) and into OB DA neurons in 10pA increments (0 – 120pA). Single AP experiments were only analysed in Spike2 if (a) they were the first AP fired in the 12-sweep 10-ms experiment; (b) the membrane potential before spiking was at -60 ± 3mV; (c) the AP peak overshot 0mV. Current threshold was defined as the current needed to elicit the first AP. Voltage threshold was measured at the point of inflexion in the depolarizing membrane potential before an AP was fired. AP properties were measured as follows: Peak at the highest voltage an AP reaches, AP afterhyperpolarization at the lowest voltage an AP reaches in the afterhyperpolarization overshoot phase after spiking; AP amplitude as the voltage difference between the voltage threshold and the AP peak of spike; AP width at half height where the half height is measured at the halfway point between the voltage threshold and the peak of spike. For OB DA neurons, the classification between biphasic/putative axon-bearing (OBb) and monophasic/putative anaxonic (OBm) DA neurons was performed based on the onset rapidness (OR) of their action potentials and visually confirmed by ML and EG. For each OB DA neuron, the first AP from their single spike experiments were extracted from Spike 2 and converted into .abf format via ABF Utility 2 (Synaptosoft Inc., NJ, USA). The truncated AP was analysed by custom-written routines in MATLAB written by Dr Matthew Grubb (Galliano et al. 2019). Voltage threshold for the analysis of onset rapidness was taken from the MATLAB algorithm as when the rate of change of membrane potential over time (dV/dt) passed 10V/s. The onset rapidness of an AP was measured from the slope of a linear fit to the phase plane plot at voltage threshold. In line with our previous findings (Chand et al. 2015; Galliano et al. 2018; 2021), DA neurons with an OR equal or larger than 5ms-1 were classified as biphasic/putative axon-bearing, DA neurons with an OR smaller than 4ms^-1^ were classified as monophasic/putative anaxonic. DA neurons with an 5*<*OR*<*4ms^-1^ are not cleanly classifiable, and were thus discarded.

For repetitive firing experiments, 500ms pulses of current of increasing amplitudes were injected at 5s intervals from 0 – 480pA at 40pA increments for midbrain DA neurons, and from 0 – 120pA at 10pA intervals for OB DA neurons. APs recorded from the multiple action potential experiments were only analysed if their membrane potential before spiking was at -60 ± 3mV and the peak of AP overshot 0mV. Input-output (I-O) firing plots were generated by plotting the number or frequency of APs against the amount of current injected in each sweep. The slope of the I-O curve measured between the sweep with a non-zero AP and the first sweep where the maximum number of APs fired throughout the experiment was recorded. The following parameters were measured only from the sweep where the maximum number of APs were fired: first action potential delay, measured by the time interval between the start of current injection (identified as when a noticeable membrane depolarization is observed) and the peak of the first AP; time between the first and last AP; average interspike interval (ms); AP frequency, measured by number of APs / time between the first and last AP * 1000.

To investigate sag potentials, 3s current injections of increasingly hyperpolarizing steps in -20pA increments were applied in current-clamp mode until the point where the initial voltage response (VP) passed -100mV. VP was the minimum voltage response within the first 100ms of the current step. To quantitatively compare sag potentials, we calculated the sag amplitude as the difference between the lowest voltage reached (VP, peak) and the steady-state (VSS, average voltage 100ms before the end of the hyperpolarization step), and the sag index (Angelo et al. 2012) using the equation [sag index = (VH – VSS)/(VH – VP)], where VH was defined as the average voltage 500ms before the current injection.

### Statistical analysis

Statistical analysis was carried out using Prism (GraphPad). “N” refers to number of animals and “n” indicates number of cells. Sample distributions were assessed for normality with the D’Agostino and Pearson omnibus test, and parametric or non-parametric tests carried out accordingly. P-values equal or smaller than 0.05 were considered significant, and all comparisons were two-tailed. For multilevel analyses, cells were nested on animals or brain areas (Aarts et al. 2014). Repetitive firing input-output curves were fitted by a mixed model with Geisser-Greenhouse correction, and Benjamini, Krieger and Yekutieli two-stage step-up method to control for false discovery rate. Principal component analysis (PCA) on electrophysiological data was performed on data standardized to have a mean of 0 and a standard deviation of 1. Principal components (PCs) were selected based on the Kaiser rule, where PCs were selected only if they had an eigenvalue greater than one. This selection was also evaluated with a scree plot. All PCAs were unsupervised; each cell was later colored on the plot by its respective location, allowing us to visually assess functionally significant clustering.

Loading scores were calculated based on standardized data and calculated using the following formula: (Eigenvector*sqrt(Eigenvalue)).

## Results

### DAT-TH colocalization rates and morphological differences between and within midbrain and forebrain DA neurons

When crossed with floxed fluorescent reporter lines, the transgenic mouse line DAT-Cre (Bäckman et al. 2006a) allows for live visualization of DA cells expressing the dopamine transporter (DAT). To assess the veracity and penetrance of DAT-tdT labelling across the dorso-ventral axis in midbrain and forebrain (Lammel et al. 2015; Papathanou et al. 2019; Stuber, Stamatakis, and Kantak 2015), we performed immunohistochemistry in fixed horizontal brain slices from juvenile animals. To enhance the genetic tomato label we used an anti-DsRed antibody and co-stained the brain slices sampled at different dorso-ventral planes with antibodies against tyrosine hydroxylase (TH), the rate-limiting enzyme in the biosynthesis of catecholamines (Figure 1A-B). In the SNC, the vast majority of cells positive for DsRed were also stained by TH, indicating excellent overlap of DAT and TH expression (double positive 99.34 ± 0.6%, DsRed+/TH-0.44 ± 0.44%, DsRed-/TH+ 0.22 ± 0.22%, n=658, N=4; Fig. 1C), a pattern of penetrance consistently high across the dorsoventral axis (Fig. 1D; Chi-squared test, p=0.95). In the VTA the expression patterns were more heterogeneous overall (double positive 85.40 ± 5.32%, DsRed+/TH-8.75 ± 4.43%, DsRed-/TH+ 6.00 ± 2.35%, n=752, N=4; Fig. 1C), and the colocalization in dorsal slices was particularly variable (Fig.1D, Chi-squared test, p=0.02). In the OB, in line with previous reports (Galliano et al. 2018; Byrne et al. 2022) only around 70% of cells expressed both the genetically-encoded DAT-tomato label and TH (double positive 72.17 ± 3.39%, DsRed+/TH-11.7 ± 1.50%, DsRed-/TH+ 16.05 ± 2.35%, n=2652, N=4; Fig. 1C). The penetrance of the genetic label was equally incomplete across the dorso-ventral axis (Fig. 1D, Chi-squared test, p=0.77) and the rostro-caudal axis (Chi-squared test, p=0.91). To match the subsequent electrophysiological experiments where only DAT-positive cells were recorded in acute brain slices, the few DsRed-/TH+ cells imaged in the fixed-tissue configuration were removed from further morphological analysis. The soma area of DAT-tdT positive neurons varied considerably within the midbrain (Fig. 1E, Table 1, nested t-test, p= 0.03), and between midbrain and forebrain (Fig. 1E, Table 1, nested t-test, p*<*0.001), with SNC neurons being 40% larger than VTA cells and almost triple the size of OB neurons. All three soma area distributions were not Gaussian with positive values for both skewness and kurtosis, but the forebrain was particularly asymmetrical. Consistent with previous work (Galliano et al. 2018; Pignatelli et al. 2005), the OB soma size distribution peaked around 70μm^2^(putative anaxonic neurons capable of regenerating throughout the animal’s lifetime) and a long tail of less numerous cells with areas overshooting 100μm^2^(putative axon-bearing neurons born exclusively during early embryonic development). Across all brain areas, the DAT-tdT neurons’ maximum soma diameter strongly correlated with its area (Fig. 1F, Table 1). Moreover, forebrain DAT-tdT cells were rounder than in midbrain as well as smaller, while SNC neurons were the largest and least round (Fig. 1G, Table 1). Finally, we found that despite the soma size differences, the DAT-tdT neuron density was similar between forebrain and midbrain (Figure 1H, Table 1, nested t-test, p=0.19), and within each brain area along the dorso-ventral axis - although density within the VTA was rather heterogeneous (Figure 1I).

**Table 1.**
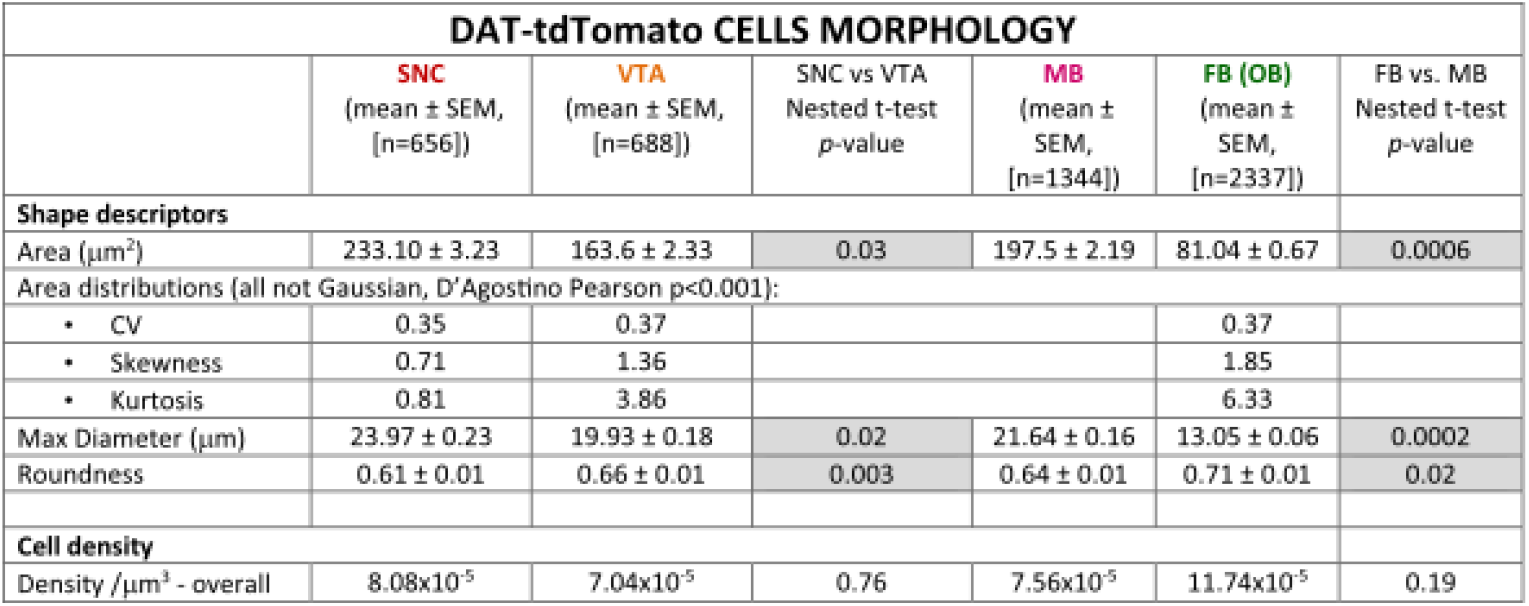
Morphological properties of DA cells. Mean values ± SEM of cell shape descriptors and cell density for DAT-tdTomato positive cells across forebrain and midbrain [N=4]. Statistical differences were calculated with a mixed model t-test nested on mouse (SNC vs VTA for within the midbrain; SNC+VTA vs OB for across brain areas). Normality of the soma area distributions was calculated with a D’Agostino Pearson Omnibus test; CV= coefficient of variation; compared with a Gaussian distribution (0 skewness and 0 kurtosis) a positive skew indicates distribution with a longer right tail and a positive kurtosis indicates a distribution with more values in the tails. Grey shading indicates statistically significant difference. Individual data points and example images are presented in Figure 1.

**Figure 1.**
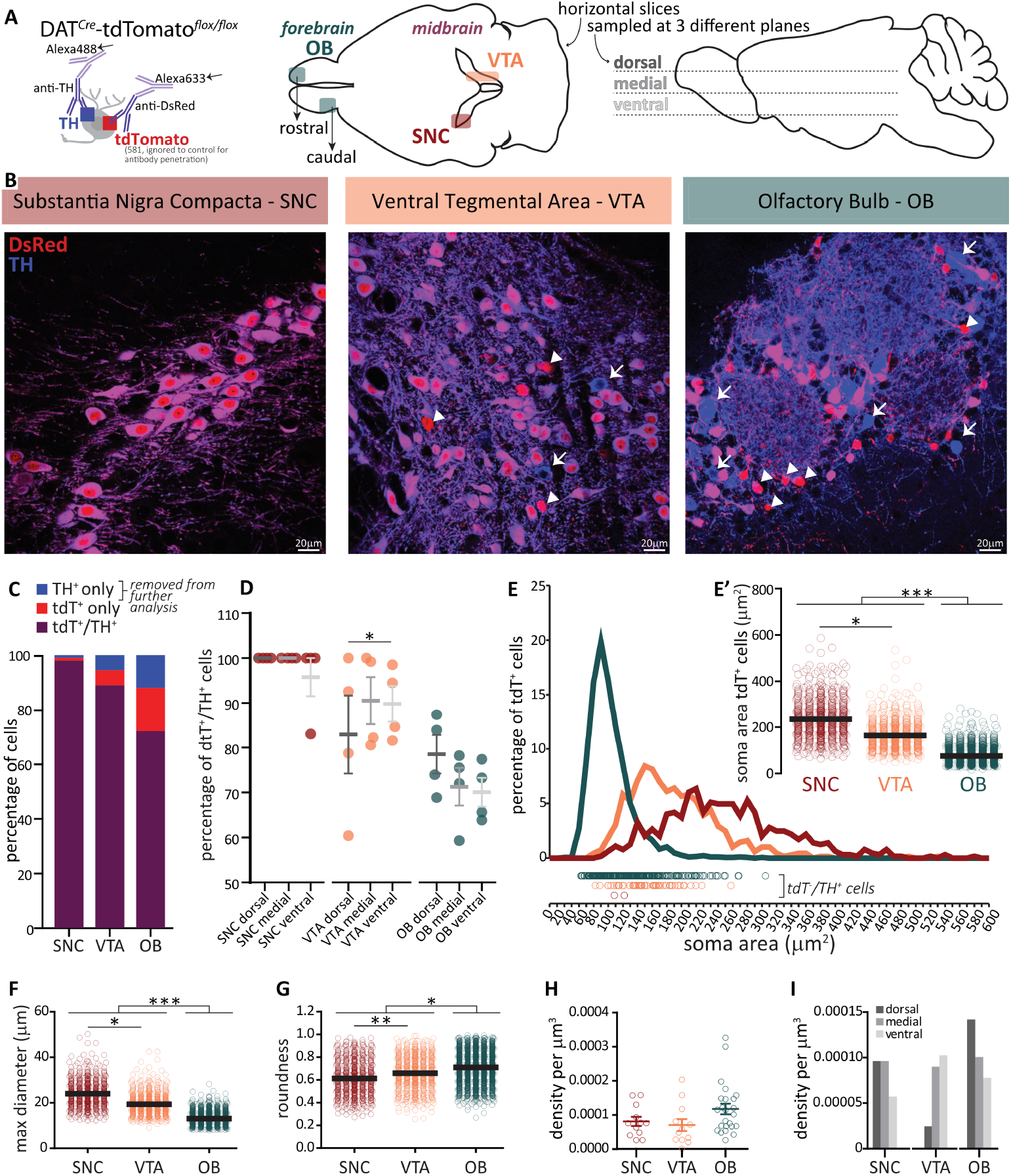
Genetic label penetrance and neuronal morphology differences in forebrain and midbrain. (A) Left: schematic representation of the immunolabelling strategy for tyrosine hydroxylase (TH) and DAT-tdTomato expression enhanced with anti-DsRed antibodies. Centre: location of forebrain olfactory bulb (OB) and midbrain substantial nigra compacta (SNC) and ventral tegmental area (VTA) DA neurons in the horizontal plane. Right: horizontal slices sampling across the dorso-ventral axis; the OB was additionally sampled across the rostro-caudal axis as indicated in the central schematic. **(B)** Example labelling of DsRed-enhanced tdTomato (red) and TH (blue) in SNC (left), VTA (centre) and OB (right). Triangles indicate cells positive for DsRed and negative for TH, and vice versa arrows indicate cells negative for DsRed and positive for TH. **(C)** Percentage of all cells labelled that were positive for both DsRed and TH (purple), DsRed only (red) or TH only (blue), in SNC (n=658, N=4), VTA (n=752, N=4) and OB (n=2652, N=4). **(D)** Percentage of cells positive for both DsRed and TH in dorsal, medial and ventral slices in the three brain areas. **(E)** DAT-tdTomato positive cells (DsRed+/TH+ and DsRed+/TH-) soma area distribution in SNC (red), VTA (orange) and OB (green); circles above x-axis indicate the area of individual DsRed+/TH-neurons. Inset in E’: individual soma area values and median for SNC (n=656, N=4), VTA (n=688, N=4), and OB (n=2337, N=4). **(F)** Maximum somatic diameter and **(G)** soma roundness of DAT-tdTomato positive neurons. **(H)** Density of DAT-tdTomato positive neurons per μm^3^ in each brain region and **(I)** in the dorsal, medial or ventral tier of each region. Circles are individual data points, thick lines are mean values, *p*<*0.05; **p*<*0.01, *p*<*0.001; for further quantification see Table 1.

### Passive electrical properties differ between the forebrain and midbrain populations of DA neurons

To investigate the functional similarities/differences between and within forebrain and midbrain DA neurons, we switched to an acute slice configuration and performed whole-cell patch-clamp recordings in DAT-tdT neurons. Given that neuronal excitability is partially determined by the passive electrical properties of the plasma membrane, we compared the neurons’ passive properties in response to a subthreshold voltage step (5mV for 10ms; Figure 2A). The resting membrane potential (Rm) was not different between midbrain and forebrain DA neurons overall, nor between the midbrain subpopulations in SNC and VTA (Figure 2B, Table 2). As reported previously (Galliano et al. 2018; 2021), with whole-cell patch clamp it is possible to further classify DA subtypes in the forebrain. Specifically, anaxonic vs axon-bearing OB DA subtypes can be teased apart on the basis of their action potential (AP) planar phase plot and onset rapidness (see methods for details). In line with previous work, we found that Rm was significantly more hyperpolarized in putative axon-bearing DA cells (OBb, with a biphasic AP phase plot) compared with putative anaxonic DA cells (OBm, with a monophasic AP planar phase plot; Fig. 2B, Table 2). Echoing the soma area differences reported in Fig. 1E, we found that membrane capacitance - a measure of the amount of plasma membrane - was overall significantly lower in forebrain than in midbrain DA neurons, once more indicating that OB neurons are smaller in size (Figure 2C, Table 2). Within the midbrain we found that SNC and VTA neuron capacitance was not significantly different, while within the forebrain the putative anaxonic OBm neurons had lower capacitance value, results in line with consistent reports of them being the smallest population (Galliano et al. 2018; Kosaka, Pignatelli, and Kosaka 2020; Chand et al. 2015; Korshunov et al. 2020) (Figure 2C, Table 2). Passive membrane resistance (Ri), a measure of the number of ion channels open at rest, was not different between the two brain areas, and there was no significant difference between SNC and VTA DA neurons. However, Ri was significantly higher in OBm than OBb neurons, indicating a less active and more electrotonically compact neuronal subtype (Figure 2D, Table 2). Together, these data indicate that OB DA neurons are smaller than midbrain DA neurons and have a higher membrane resistance, but a similar resting membrane potential.

**Table 2.** Electrophysiological properties of DA cells. Mean values ± SEM of passive properties, sag currents, action potential properties (SNC n=16, VTA n=16, OBb n=14, OBm n=23) and repetitive firing properties (SNC n=13, VTA n=14, OBb n=14, OBm n=15) for DA cells across forebrain and midbrain. Statistical differences between brain areas (forebrain vs. midbrain) were calculated with a mixed model t-test nested on sub-region. Within each brain region, differences between sub-regions groups were calculated independently with an unpaired t-test for normally-distributed data (“t” for data with equal variance, “tW” for data with unequal variance and Welch correction) or with a Mann–Whitney test for non-normally distributed data (“MW”). Grey shading indicates statistically significant difference. Individual data points and example traces are presented in Figures 2-5.

| DAT-tdTomato CELLS PHYSIOLOGY |  |  |  |  |  |  |  |  |  |
| --- | --- | --- | --- | --- | --- | --- | --- | --- | --- |
|  | MB<br>mean<br>± SEM | FB<br>mean<br>± SEM | MB vs FB<br>Nested<br>t-test<br>*p*-value | SNC<br>mean<br>± SEM | VTA<br>mean<br>± SEM | SNC vs<br>VTA<br>Test type,<br>*p*-value | OBb<br>mean<br>± SEM | OBm<br>mean<br>± SEM | OBb vs<br>OBm<br>Test type,<br>*p*-value |
| **PASSIVE PROPERTIES** |  |  |  |  |  |  |  |  |  |
| Resting potential (mV) | -46.25 ± 0.91 | -50.03 ± 0.96 | 0.15 | -45.56 ± 0.99 | -46.94 ± 1.54 | t, 0.46 | -52.86 ± 1.49 | -48.30 ± 1.13 | t, 0.02 |
| Membrane capacitance (pF) | 55.82 ± 2.53 | 26.35 ± 1.57 | 0.04 | 59.31 ± 3.25 | 52.33 ± 3.79 | t, 0.17 | 31.85 ± 2.62 | 23.00 ± 1.64 | MW, 0.01 |
| Input Resistance (mΩ) | 194 ± 13 | 420 ± 35 | 0.15 | 184 ± 20 | 204 ± 17 | MW, 0.19 | 310 ± 43 | 488 ± 46 | MW, 0.01 |
| **SAG VOLTAGE** |  |  |  |  |  |  |  |  |  |
| Min voltage (peak, mV) | -105 ± 0.7 | -106 ± 1.2 | 0.78 | -103 ± 0.9 | -106 ± 1.0 | MW, 0.14 | -103 ± 1.5 | -107 ± 1.5 | MW, 0.01 |
| Steady state (mV) | -86 ± 1.6 | -100 ± 0.8 | 0.06 | -82 ± 1.4 | -90 ± 2.5 | t, 0.02 | -99 ± 1.3 | -101 ± 1.1 | MW, 0.21 |
| Sag amplitude (mV) | 18.71 ± 1.27 | 5.49 ± 0.67 | 0.04 | 21.39 ± 0.98 | 16.03 ± 2.17 | MW, 0.07 | 4.05 ± 0.58 | 6.37 ± 0.98 | MW, 0.15 |
| Sag index | 0.58 ± 0.03 | 0.89 ± 0.02 | 0.046 | 0.51 ± 0.03 | 0.65 ± 0.05 | tW, 0.02 | 0.91 ± 0.93 | 0.87 ± 0.02 | tW, 0.11 |
| **ACTION POTENTIAL PROPERTIES** |  |  |  |  |  |  |  |  |  |
| Threshold (pA/pF) | 11.44 ± 1.14 | 3.23 ± 0.21 | 0.15 | 7.80 ± 0.65 | 15.09 ± 1.79 | tW, 0.001 | 2.37 ± 0.37 | 3.75 ± 0.18 | tW, 0.004 |
| Threshold (mV) | -35.85 ± 1.13 | -36.65 ± 0.54 | 0.64 | -37.04 ± 1.44 | -34.66 ± 1.74 | MW, 0.30 | -38.34 ± 0.99 | -35.61 ± 0.52 | t, 0.01 |
| Onset rapidness (1/ms) | 17.69 ± 1.06 | 5.05 ± 0.56 | 0.05 | 17.77 ± 1.70 | 17.60 ± 1.34 | t, 0.93 | 8.595 ± 0.81 | 2.88 ± 0.18 | n/a |
| Peak amplitude (mV) | 33.81 ± 2.08 | 47.56 ± 1.08 | 0.09 | 29.46 ± 3.13 | 38.17 ± 2.38 | t, 0.03 | 47.64 ± 1.74 | 47.51 ± 1.41 | t, 0.96 |
| Width at half-height (ms) | 0.78 ± 0.03 | 0.76 ± 0.02 | 0.68 | 0.76 ± 0.04 | 0.79 ± 0.04 | t, 0.60 | 0.69 ± 0.03 | 0.80 ± 0.03 | t, 0.03 |
| After hyperpolarization (AHP, mV) | -27.98 ± 1.11 | -39.67 ± 0.87 | 0.06 | -25.37 ± 1.26 | -30.58 ± 1.61 | t, 0.02 | -38.15 ± 1.60 | -40.60 ± 0.98 | t, 0.17 |
| **REPETITIVE FIRING PROPERTIES** |  |  |  |  |  |  |  |  |  |
| Max number of APs | 7.63 ± 1.04 | 25.17 ± 2.01 | 0.03 | 5.69 ± 0.41 | 9.43 ± 1.87 | MW, 0.02 | 27.79 ± 2.74 | 22.73 ± 2.87 | t, 0.21 |
| First AP delay (ms) | 267 ± 22 | 109 ± 21 | 0.15 | 202 ± 19 | 327 ± 30 | t, 0.002 | 136 ± 34 | 83 ± 23 | MW, 0.14 |
| Inter-spike interval CV | 0.47 ± 0.04 | 0.19 ± 0.02 | 0.05 | 0.53 ± 0.06 | 0.42 ± 0.06 | t, 0.17 | 0.15 ± 0.02 | 0.22 ± 0.04 | MW, 0.11 |
| Max AP frequency (Hz) | 51.59 ± 5.53 | 75.38 ± 5.41 | 0.29 | 66.85 ± 8.44 | 37.42 ± 4.98 | t, 0.01 | 67.76 ± 7.01 | 82.50 ± 7.95 | t, 0.18 |
| Slope I/O curve (Hz/(pA/pF)) | 3.17 ± 0.85 | 7.94 ± 0.51 | 0.08 | 4.61 ± 1.66 | 1.82 ± 0.38 | MW, 0.02 | 8.14 ± 0.77 | 7.75 ± 0.70 | t, 0.71 |

**Figure 2.**
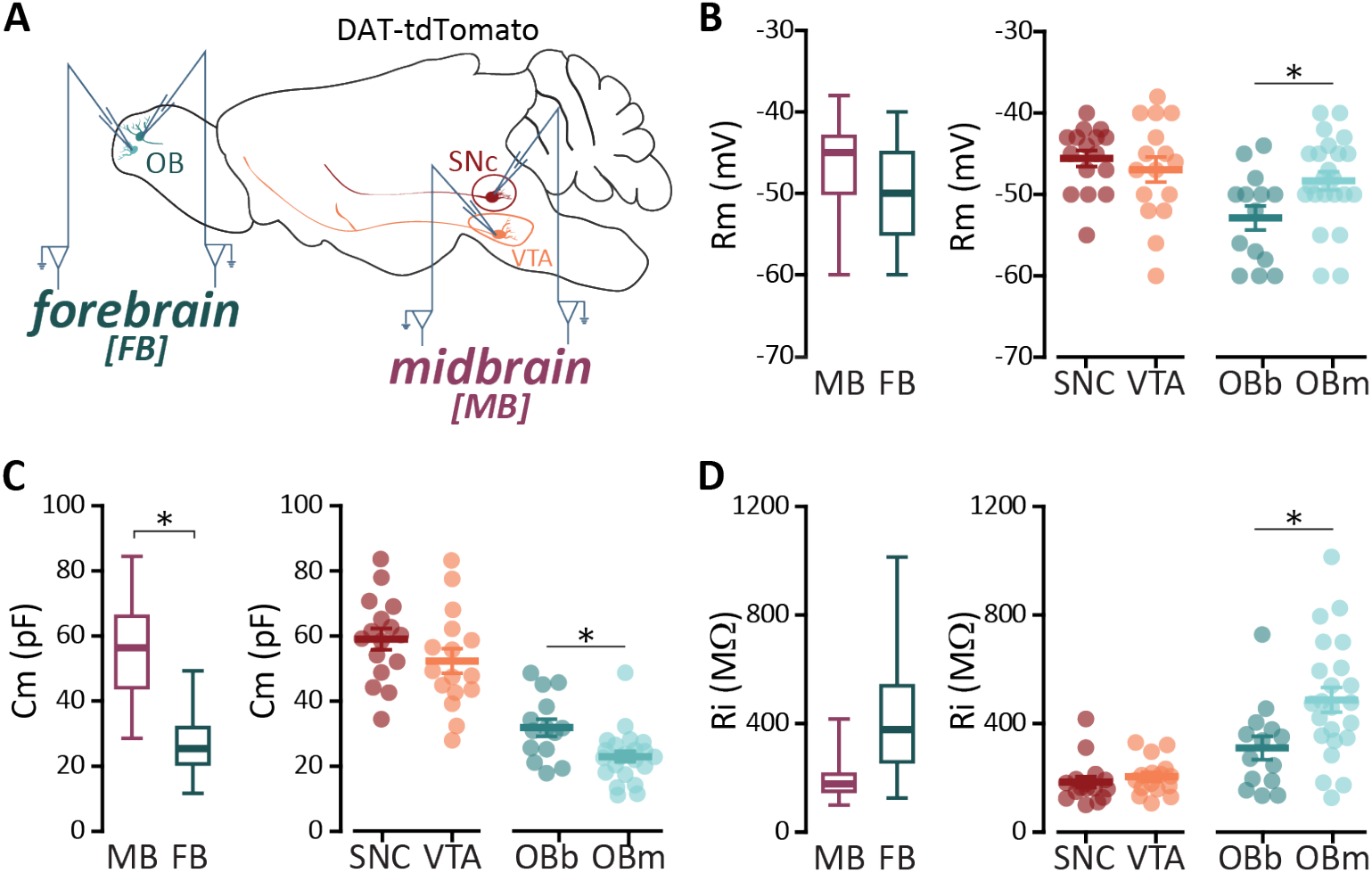
Passive electrical properties differ between forebrain and midbrain, and between the two subpopulations of OB DA neurons. **(A)** Schematic representation of the location of OB DA neurons in the forebrain and VTA and SNC DA neurons in the midbrain, targeted for whole cell patch clamp recordings in acute horizontal brain slices from juvenile DAT-tdTomato mice. **(B)** Resting membrane potential (Rm) in midbrain (SNC n=16, VTA n=16) and forebrain (OBb n=14, OBm n=23) neurons overall (left) and divided by subtype (right). **(C-D)**. As in B, for membrane capacitance (Cm) and input resistance (Ri). Circles are individual cells, lines are mean ± SEM, *p*<*0.05; further quantification and statistical analysis in Table 2.

### Voltage sag responses to hyperpolarization are stronger in mid-brain than forebrain DA neurons

In the midbrain, identification of DA neurons is often determined in voltage clamp by the presence of a hyperpolarization-activated cation current (Ih, or current sag) which plays a pivotal role in controlling intrinsic excitability (Neuhoff et al. 2002). In current clamp recordings, the time- and voltage-dependent opening of HCN channels results in a slow depolarizing response (steady-state) to a triggered hyperpolarization (peak), referred to as a voltage sag. By injecting hyperpolarizing current steps in the four DA groups, we found no differences between forebrain and midbrain in the raw measurement of sag minimum voltage peak, hyperpolarization amplitude, and steady state. However, when we computed the voltage sag amplitude as the difference between steady-state and peak, and sag index as the ratio between steady-state and peak voltage corrected for holding voltage (Angelo et al. 2012) (Fig. 3B), we found, respectively, higher and lower values in midbrain DA neurons than in forebrain cells (Figure 3C-D; Table 2). In line with previous data from rats (Pignatelli et al. 2013), the two subtypes of murine OB neurons displayed similarly shallow voltage sags, or no sag at all. Conversely, within the midbrain we found higher sag indexes – and a trend towards higher sag amplitudes – in more SNC than VTA cells, which once more displayed more heterogeneity (Figure 3C-D; Table 2; sag amplitude coefficient of variation [CV] in SNC=0.18, VTA=0.54). Finally, we quantified the percentage of cells firing rebound action potentials following a hyperpolarization-induced voltage sag. While approximately half of all OB DA neurons fired on rebound (57% of OBb, 48% of OBm), almost no midbrain DA neuron generated an action potential following the end of the hyperpolarizing step (SNC 6%, VTA 0%; 2 test, p*<*0.001; Figure 3E). Taken together, these data show that SNC midbrain DA neurons have the largest voltage sag, followed by VTA DA neurons. Conversely, OB neurons of both categories have small voltage sags but are likely to fire rebound action potentials following a sustained hyperpolarization.

**Figure 3.**
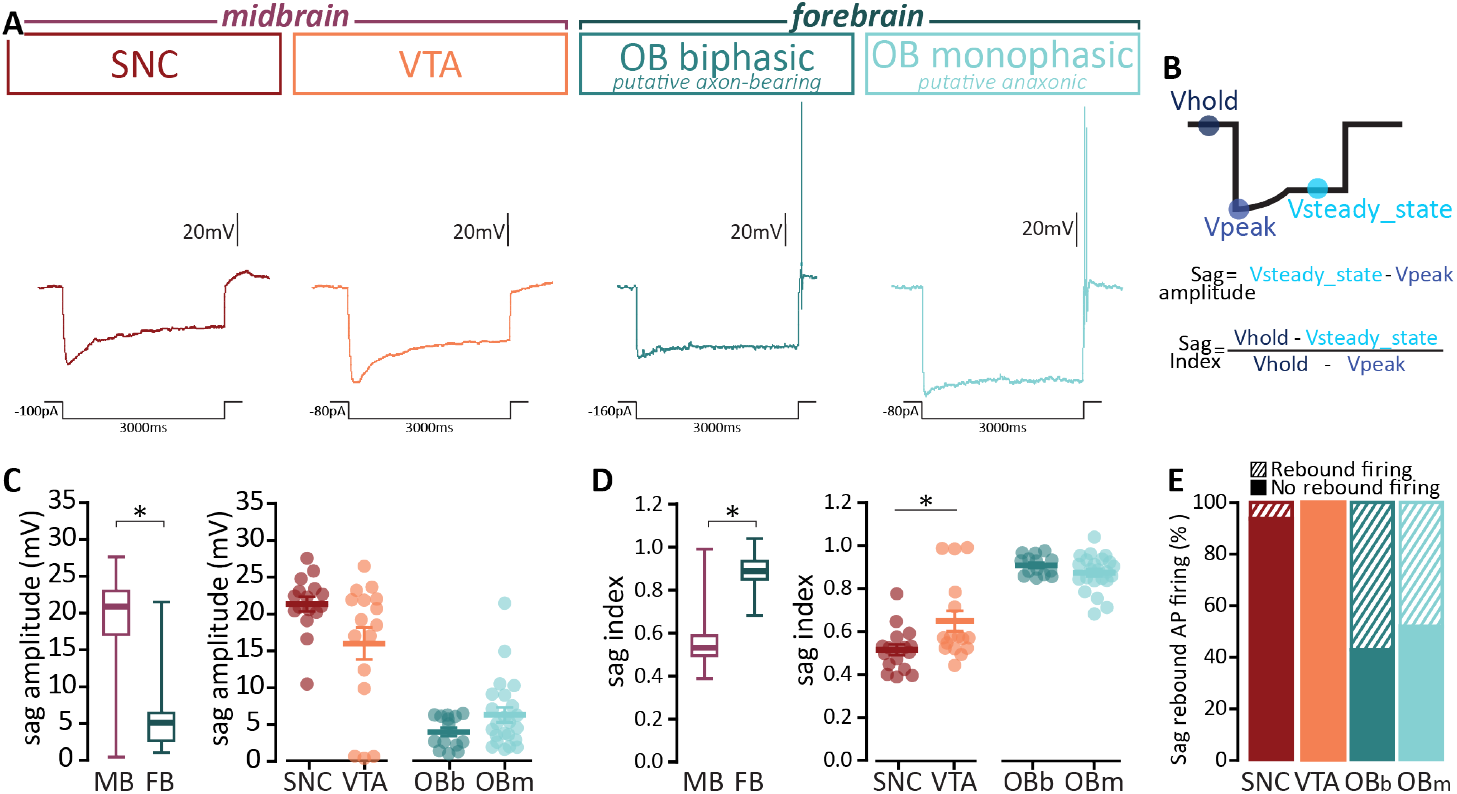
The depolarization response to hyperpolarization (sag potential) is consistently present in midbrain DA neurons, while it is small or absent in forebrain DA cells. **(A)** Example traces of the voltage sag response to hyperpolarizing current injections in midbrain (SNC n=16, VTA n=16; left) and forebrain (OBb n=14, OBm n=23; right) DA neurons. **(B)** Schematic visualization of the voltage sag analysis parameters and formulas used to calculate sag amplitude and index. **(C)** Voltage sag amplitude and **(D)** sag index in midbrain and forebrain neurons overall (left) and divided by subset (right). **(E)** Percentage of neurons firing an action potential (striped bars) after the rebound depolarization following hyperpolarization. Circles are individual cells, lines are mean ± SEM, *p*<*0.05; further quantification and statistical analysis in Table 2.

### Action potential waveforms are similar in forebrain and midbrain DA neurons

To analyse the properties of individual action potentials fired at threshold, we injected 10ms depolarizing current steps of increasing amplitude in the four groups of DA neurons (Figure 4A). Individual action potentials in forebrain and midbrain DA neurons were remarkably similar, showing no overall differences in voltage and injected current density threshold, and width at half-height (Fig. 4B,C,F, Table 2). The lack of difference between forebrain and midbrain for action potential current threshold might be explained by the large variance in the data obtained from VTA DA neurons (interquartile range SNC= 3.6pA/pF, VTA=10.2pA/pF, OBb=2.2pA/pF, OBm=1.2pA/pF; Figure 4B), likely reflecting the heterogeneity of this population. Albeit not statistically different, the AP onset rapidness, which we used to classify the OB subtypes given the characteristically low values in the OBm cells lacking the axon initial segment initiation site, trended towards higher values in the midbrain (Fig. 4D, p=0.05). Likewise, AP and after-hyperpolarization (AHP) peak amplitudes exhibited strong trends towards higher, but not statistically different, values in the forebrain (Fig. 4E,G, Table 2). When we analysed differences within each brain area, we confirmed that the putative axon-bearing OBb DA cells require less current and a less depolarized voltage than their putative-anaxonic OBm counterparts to fire an action potential (Fig. 4B-G, Table 2). Within the midbrain, we found marked differences in terms of firing threshold (half the injected current density needed in SNC vs. VTA cells), and AP shape with lower peak and AHP amplitudes in SNC than VTA (Fig. 4B-G, Table 2). Overall, these data suggest that a similar but not identical complement of voltage gated ion changes underlie the action potential in all four neuronal types. Notably, all four classes of DA neurons have large amplitude AHPs (Figure 4G).

**Figure 4.**
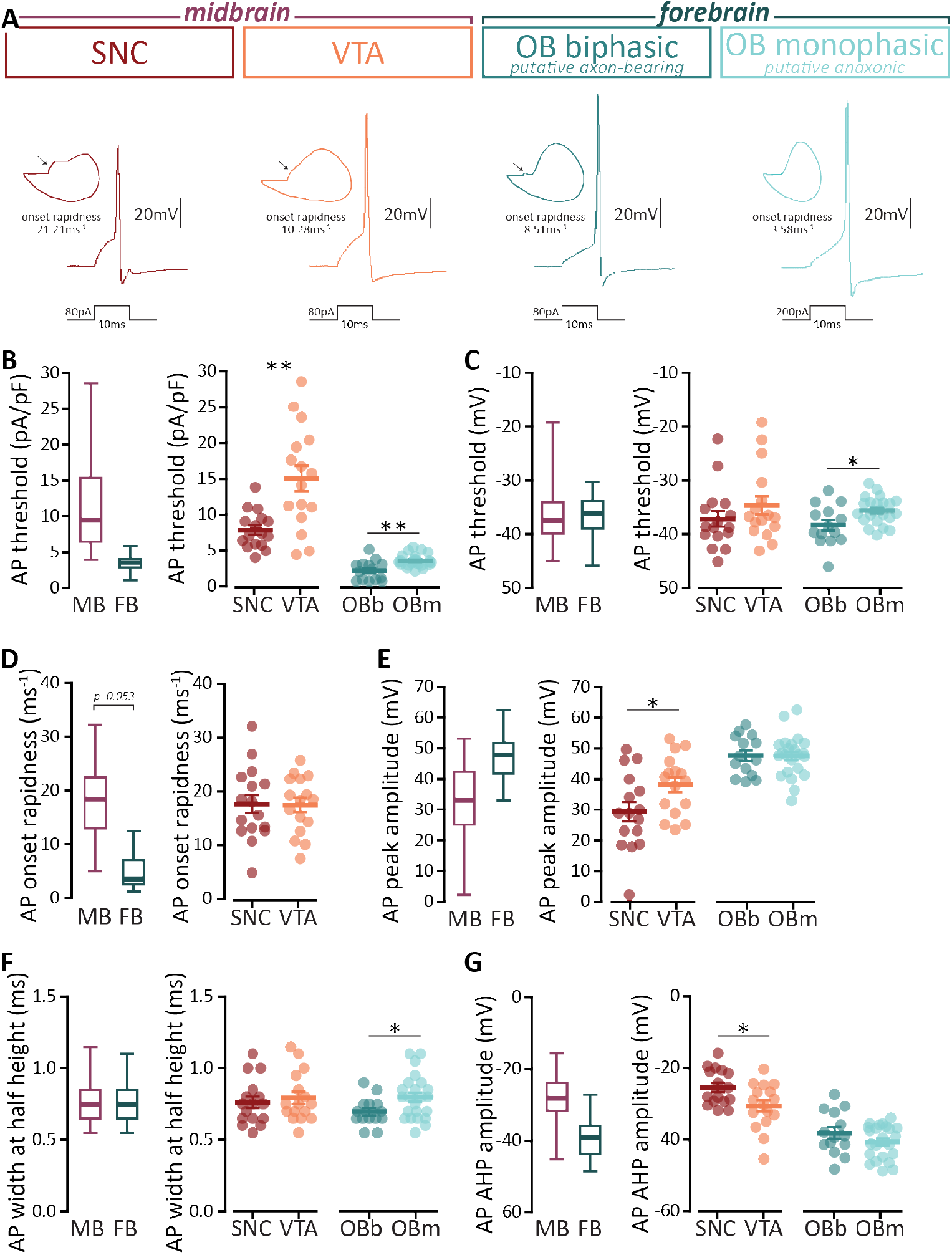
Action potential waveforms are largely similar across forebrain and midbrain DA neurons, but vary substantially between SNC and VTA. **(A)** Example traces of the membrane voltage response to the minimum depolarizing 10ms-long current injection needed to evoke an AP in midbrain (SNC n=16, VTA n=16; left) and forebrain (OBb n=14, OBm n=23; right) DA neurons. Arrow on the planar phase plots points to the axon initial segment-dependent first AP phase. Waveform parameters for midbrain and forebrain neurons overall (left) and divided by subset (right) include **(B)** injected current density needed to evoke an AP, **(C)** membrane potential at which the AP was evoked, **(D)** AP rate of rise from threshold (onset rapidness), **(E)** AP peak membrane potential, **(F)** AP width at half the maximum height, and **(G)** peak amplitude of the AP after hyperpolarization (AHP). Circles are individual cells, lines are mean ± SEM, *p*<*0.05; **p*<*0.01, further quantification and statistical analysis in Table 2.

### Repetitive action potential firing differs between forebrain and midbrain

Despite the similarity in their AP waveforms, forebrain and midbrain DA neurons displayed remarkably different repetitive firing properties when probed with 500ms-long current injections of increasing intensity (Fig. 5A-E). To allow for inter-area comparisons despite the substantial differences in cell size, we normalized current injections for cell capacitance (*i*.*e*., current density) and confirmed different input-output curves between the midbrain and forebrain DA neurons, and at high current densities between the two OB subtype (Fig. 5F, mixed-model ANOVA, effect of manipulation F1.38, 56 = 82.16, p*<*0.001, effect of time F3, 52 = 28.69, p*<*0.001; effect manipulation x time F27, 376 = 14.36, p*<*0.001; OBb vs OBm for injected current*>*5.5pA/pF, p*<*0.05). The most striking difference between forebrain and midbrain DA cells’ repetitive firing was the maximum number of APs fired within the injection window, which was over three times larger in the forebrain than in the midbrain (Figure 5G, Table 2). While the timing of the first AP is squarely similar in forebrain and midbrain DA neurons (Fig. 5H), the inter-spike interval coefficient of variation (ISI CV) was lower for the forebrain (Fig. 5I, p=0.048). The overall higher excitability in the forebrain as measured by maximum number of APs was mirrored in a strong trend towards an higher slope of the current density vs. AP frequency input-output curve (Fig. 5J-K, Table 2, p=0.08). While we only found differences between the two bulbar DA subtypes at high current injections (but reported strong trends towards OBb being overall more excitable), we observed that SNC and VTA differ in almost all parameters (Fig. 5G-K, Table 2). Of note once more is the variability observed in VTA: for instance, while most midbrain neurons fired less than 10 action potentials, one VTA neuron fired a maximum of 30 spikes (interquartile range SNC=1.5, VTA=4; coefficient of variation SNC=0.26, VTA=0.74).

**Figure 5.**
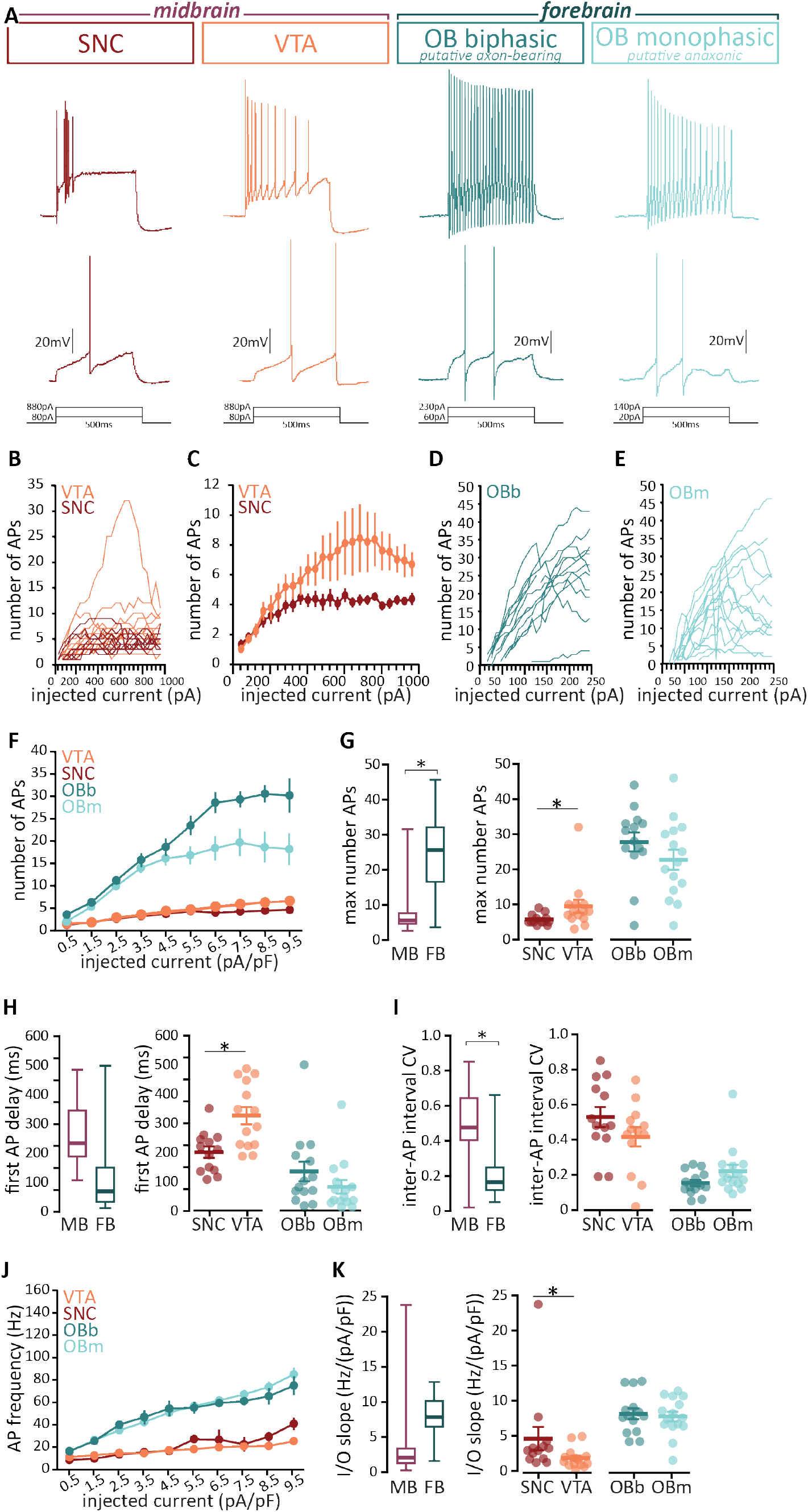
Repetitive action potential firing differ between forebrain and midbrain DA neurons, and between SNC and VTA cells. **(A)** Example traces of the membrane voltage response to 500ms depolarizing current injections of increasing intensity evoking repetitive AP firing, in midbrain (SNC n=13, VTA n=14; left) and forebrain (OBb n=14, OBm n=15; right) DA neurons. **(B)** Input-output plots showing the number of APs fired at each current injection in individual SNC (red) and VTA (orange) DA neurons. **(C)** Mean number of APs and SEM at each current injection in SNC and VTA neurons. **(D**,**E)** As in (B), input-output plots for individual OBb (teal) and OBm (turquoise) DA neurons. **(F)** Mean number of APs and SEM at each current density (*i*.*e*., injected current normalized for cell capacitance) in SNC (red), VTA (orange), OBb (teal) and OBm (turquoise) DA neurons. Repetitive firing parameters for midbrain and forebrain neurons overall (left) and divided by subset (right) include **(G)** maximum number of APs, **(H)** latency of the first AP at the current injection level where the max AP number was fired, (**I)** coefficient of variation of the inter-spike intervals (ISI CV) at the current injection level where the max AP number was fired. **(J)** Mean AP firing frequency and SEM plotted against injected current density and (K) slope of these frequency vs. current density input-output curves for midbrain and forebrain neurons overall (left) and divided by subset (right). Circles and thin lines are individual cells, thick lines are mean ± SEM, *p*<*0.05; further quantification and statistical analysis in Table 2.

### Principal component analysis highlights consistent grouping of midbrain vs forebrain subtypes

Finally, to understand the overall impact of the various individual electrophysiological features reported above, we performed principal component analyses (PCA; see Materials and Methods for details). The first two principal components (PC) accounted for 43.16% and 13.94% of the variance respectively (Fig. 6A-B). Plotting the primary and secondary component scores for each neuron against each other revealed clear clustering of forebrain and midbrain DA neurons (Fig. 6A). To determine whether a specific set of electrophysiological properties predominantly contributed to PC1 and PC2, we graphed loading scores (derived from eigenvalues and eigenvectors) for each PC and found no clear-cut major single source of variation (Fig. 6C). To further confirm that no single experimental protocol – passive properties, voltage sag, AP shape, repetitive firing – alone drove the variability in the dataset, we performed two additional PCAs. The first included only measurements derived from single and repetitive AP firing protocols and once more returned two clear midbrain vs. forebrain clusters with PC1 and PC2 responsible for 42.64% and 17.82% of the variation respectively (Fig. 6D). Similarly, PCA analysis of sag voltage measurements resulted in similar but clear clustering patterns (PC1: 63.30%; PC2: 33.60%; Fig. 6E). Across all three PC analyses we found some overlap between the midbrain and forebrain clusters (1-3 “misplaced” cells). This is perhaps unsurprising given the variability of electrophysiological properties, especially in VTA. In sum, although we found considerable overlap in individual intrinsic properties of DA neurons across the two brain areas, when all measurements are evaluated holistically we found a clear functional split between forebrain and midbrain DA neurons.

**Figure 6.**
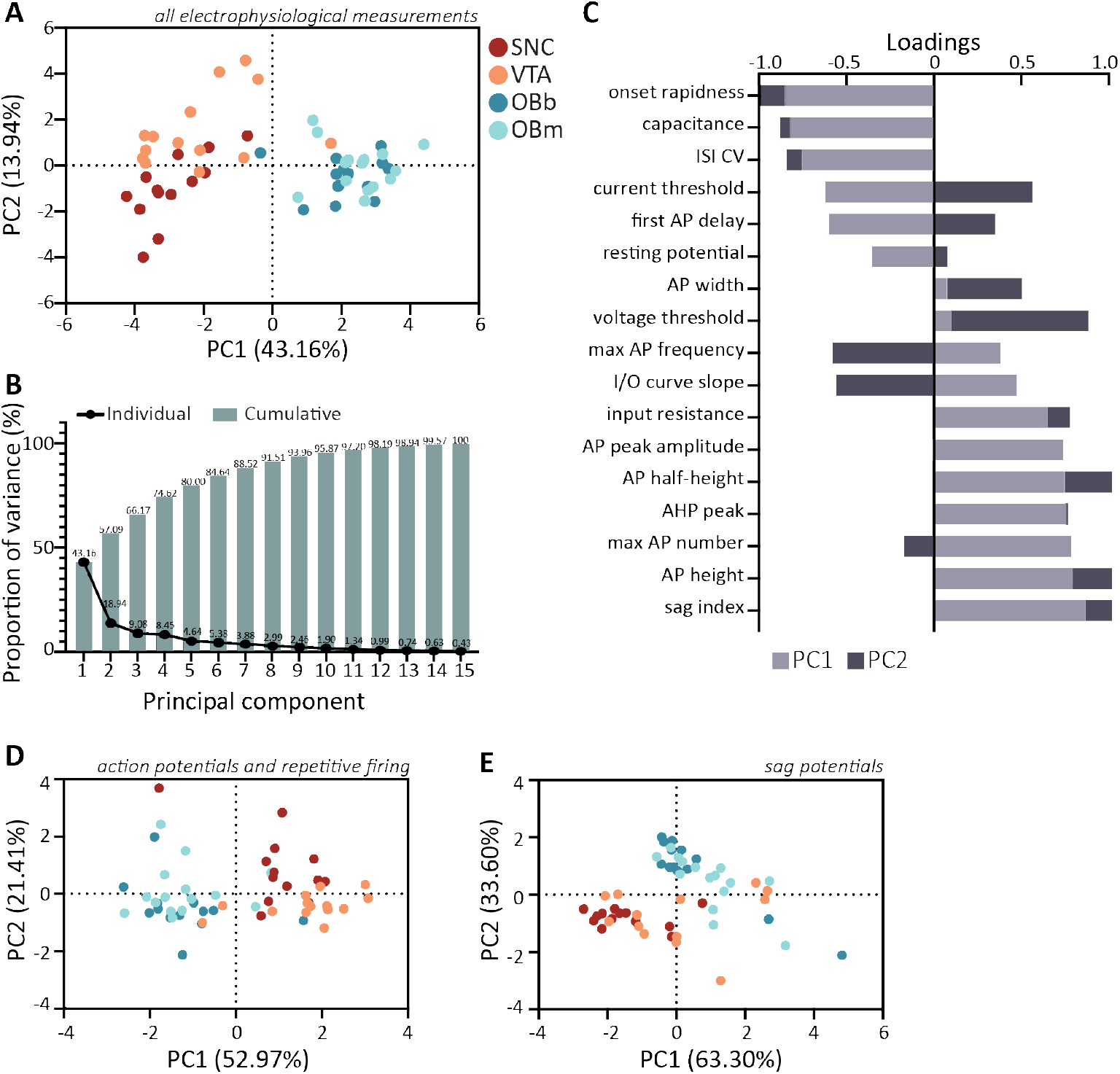
Principal component analysis (PCA) of electrophysiological properties reveals clear clustering of midbrain and forebrain DA neurons. **(A)** PC score plot for forebrain (OBb, teal, n=14; OBm, turquoise, n=15) and midbrain (SNC, red, n=13; VTA, orange, n=14) DA neurons based on all measurements obtained from whole-cell recordings (Figures 2-5, Table 2). Each circle represents a cell plotted against its primary and secondary principal component (PC) scores. **(B)** Individual and cumulative proportion of variance explained by each principal component. **(C)** Loading scores for all variables showing their respective contributions to PC1 (light grey) and PC2 (dark grey). **(D-E)** As in C, but for PCA including only measurements from single AP + repetitive firing (Figures 4-5) and sag potentials (Figure 3), respectively.

## Discussion

In this study we compared the morphological and functional properties of midbrain and forebrain DA neurons identified via the expression of the fluorescent marker tdTomato under the control of the DAT promoter (Bäckman et al. 2006b). We demonstrated that within the midbrain, VTA and SNC DA neurons are only broadly similar in their action potential waveform and repetitive firing properties, and that VTA DA neurons showed greater heterogeneity. We also confirmed previous findings that putative anaxonic/regenerating OB DA neurons are smaller and less excitable than their embryonic-born/axon-bearing counterparts (Galliano et al. 2018). Interestingly midbrain and forebrain DA neurons were hugely different in terms of sag voltage and excitability, indicating that DA neurons do not share a common neurophysiological signature.

### Penetrance and specificity of the DAT-Cre line

Given that, to the best of our knowledge, this is the first study to use the DAT-Cre line to electrophysiologically profile midbrain DA neurons, we first characterized this reporter line against the gold standard of TH expression. Previous work showed incomplete penetrance and imperfect specificity in the OB (Galliano et al. 2018; Byrne et al. 2022), a finding that we replicated: 16% of TH positive neurons failed to express DAT, and 12% of DAT-expressing neurons did not express TH (a population that has been previously characterized as calretinin-positive and whose electrophysiological signatures clearly distinguishes them from DA cells (Fogli Iseppe, Pignatelli, and Belluzzi 2016; Byrne et al. 2022)). Conversely, in SNC and VTA over 90% of DAT-tdTomato neurons co-stained for anti-TH antibodies, indicating that the DAT-Cre line captures most midbrain DA neurons with minimal leakage. One could speculate that such a discrepancy in DA neuron coverage between midbrain and forebrain with the DAT-Cre line can be caused by more abundant polymorphisms in the Slc6a3 gene and/or different allele dosage, as well as by the ectopic expression of the gene in the bulbar calretinin interneurons (Stuber, Stamatakis, and Kantak 2015).

### Morphological and functional heterogeneity of midbrain DA neu-rons

A plethora of circuit functions and behavioural roles has been attributed to midbrain DA neurons (Berke 2018), and it is now accepted that such range at the systems level is matched by a similar breadth at the molecular and cellular level (Garritsen et al. 2023). *In vivo*, synaptic inputs enable midbrain DA neurons to fire action potentials in bursting and irregular patterns (Grace and Bunney 1984), but the strong influence of intrinsic ion channels imposes a more regular firing pattern *ex vivo* (Gantz et al. 2018). Across the various protocols assessing passive and active electrophysiological properties, our DAT-tdTomato mouse SNC recordings compare well with previously published rat data (Pucak and Grace 1996). Similarly, the VTA results from genetically-tagged mouse DA neurons are broadly in line with recordings of rat VTA DA neurons defined by TH and Ih expression(Margolis et al. 2006), including a similar soma size and passive properties, as well as comparable action potential waveforms. With the current step protocol that we employed, VTA DA neurons fired a maximum of 9 spikes at a maximum frequency of 37 Hz, higher than spontaneous spike firing frequency (1-10 Hz; (Margolis et al. 2006; Ungless and Grace 2012)). Given that the majority of our VTA recordings were in the ventral tier, where neurons are known to fire at higher frequencies than their dorsal counterparts (Lammel et al. 2008), such discrepancy can be explained by a sampling difference as well as by the different evoked vs. spontaneous firing configuration. Importantly, our *ex vivo* results are consistent with the high frequency phasic firing seen during reward prediction error learning (Schultz 2016) and with the heterogeneity extensively reported in the literature (Morales and Margolis 2017). Intra-group heterogeneity notwithstanding, we found substantial differences when we compared SNC vs. VTA neurons in DAT-tdTomato mice. Overall, our data show that SNC DA neurons have a larger and more oblong soma but are nevertheless more excitable than VTA DA cells when the injected current was normalized for cell capacitance. Compared to VTA neurons, SNC APs were fired at lower thresholds but had a smaller peak and AHP amplitude, and the AP number plateaued rapidly when the neurons were probed with prolonged current injections. Taken together with the consistently larger voltage sag in SNC DA cells than in the VTA neurons, our results indicate that the two midbrain subtypes are equipped with a different complement of voltage-gated channels, and that within each subgroup such complement is varied and allows for the observed heterogeneity of both action potential and sag voltage properties.

### Midbrain versus forebrain DA neurons

Forebrain DA neurons are a handful of cells which makes up a very small fraction of both bulbar inhibitory interneurons and DA neurons overall – yet, even in their scarcity, they display extreme diversity in terms of developmental profile, shape, and function (Galliano et al. 2018; Chand et al. 2015; Galliano et al. 2021; Kosaka, Pignatelli, and Kosaka 2020; Kiyokage et al. 2010; Bonzano et al. 2016). Our results confirm that using the AP rate of rise and bi-or mono-phasic nature of the planar phase plot allows for a clear split between putative-anaxonic and putative-axon bearing, with the latter being larger and more excitable. To compare the repetitive firing of large but less excitable midbrain DA neurons with the smaller but more excitable forebrain ones, we skewed our injected current steps to higher values than in our previous studies (Galliano et al. 2018; 2021). While this dataset thus lacks the granularity to finely sample firing behaviour around threshold, it confirmed that at high injected current densities OBb DA neurons had a faster-rising input-output curve than OBm cells, in line with their overall higher excitability. Interestingly, the two OB subtypes have similarly shallow voltage sags, with values at times so low that they cease to be sags altogether (Pignatelli et al. 2013). This is a very interesting difference between forebrain and midbrain DA cells, since the presence of a voltage sag and and Ih current have long been considered a signature of the dopaminergic phenotype, so much so that it was used to identify them before the advent of transgenic mouse lines. Furthermore, while forebrain and midbrain DA cells have largely similar AP waveforms, their repetitive firing behaviour is substantially different. Specifically, forebrain DA neurons of either subtype fired over thrice the number of APs than midbrain DA cells over a 500ms current injection window and did so with a much more regular temporal profile. In line with data from hippocampal pyramidal neurons (Hewitt, Ordemann, and Brager 2021), one could hypothesize that the more prominent sag in the midbrain gears the DA neurons towards a faster-adapting repetitive firing. This could be neuroprotective especially for those cells located in the ventral midbrain which co-transmit glutamate, and perhaps unneeded in the GABA-co-transmitting OB DA cells (Vaaga, Borisovska, and Westbrook 2014; Wallace and Sabatini 2023).

Our PCA analysis unexpectedly showed that overall the large, embryonic-born, axon-bearing bulbar DA cells are less similar to large, embryonic-born, axon-bearing midbrain DA cells than to their small, regenerating, anaxonic bulbar counterparts. Indeed, the striking heterogeneity that we described within each subgroup and each brain area is nonetheless overruled by the even larger differences in size, shape, and electrophysiological properties of DA neurons between forebrain and midbrain. Such findings highlight the challenges faced by cell replacement therapies aimed at harvesting the subventricular zone precursor of the bulbar DA cells and transplanting them in Parkinsonian animals (Cave, Wang, and Baker 2014). Regenerating OB neurons are anaxonic and can therefore be only transplanted directly into the striatum, not in the SNC. Moreover, even if they could manage to complete their differentiation and connect within the striatal network, they would likely fire more APs than SNC neurons, and in doing so co-release GABA as well as dopamine. Therefore, their net effect on striatal processing may not be suitable to replace the lost SNC DA drive.

### Not all dopaminergic neurons are created equal

Is there a common set of morphological and/or functional properties that makes a neuron dopaminergic? It does not seem likely. Research investigating all DA groups within the brain has been steadily accumulating, with each new study revealing an increasing level of heterogeneity. While in this study we did not record from the retina and hypothalamic DA groups, others have highlighted how those neurons too are molecularly, morphologically, and functionally diverse (Zhang, Zhou, and McMahon 2007; Hirasawa, Contini, and Raviola 2015; Romanov et al. 2017; Korchynska et al. 2022). Moreover, DA heterogeneity seems to be an evolutionarily conserved feature across phyla, with reports in both other vertebrates such as the zebrafish (Caldwell et al. 2019; McLean and Fetcho 2004) and invertebrates such as drosophila (Ma et al. 2023; Otto et al. 2020). So, what defines a DA neuron? Perhaps we ought to consider DA neurons just as varied as “typical” neurons (Tasic et al. 2016; Cerminara et al. 2015; Morgan et al. 2016). However, albeit diverse, they share a common feature: dopamine. This allows them to exert an enhanced level of temporal and spatial control on their postsynaptic targets, thereby supporting an extensive array of behavioural functions.

## Author Contributions

MYHL: Investigation, Validation, Formal analysis, Data curation.

SG: Validation, Formal analysis, Data curation, Visualization, Writing – review and editing

SJ: Conceptualization, Data curation, Formal analysis, Funding acquisition, Project administration, Supervision, Writing – original draft, Writing – review and editing

EG: Conceptualization, Investigation, Data curation, Formal analysis, Visualization, Funding acquisition, Project administration, Supervision, Writing – original draft, Writing – review and editing The authors declare no financial or personal relationship which could be construed as a potential conflict of interest.

## Acknowledgments

This work was supported by a Royal Society Research RGS-R1-191481 and an ISSF Grant to EG, and two Physiology, Development and Neuroscience Departmental Research Grants to SJ and EG. We wish to thank Matthew Grubb for antibodies and access to his lab’s confocal microscope, Ailie McWhinnie for comments on the manuscript, and all members of the Galliano laboratory for providing helpful discussions.

